# Deep spatial proteomic exploration of severe COVID-19-related pulmonary injury in post-mortem specimens

**DOI:** 10.1101/2023.07.14.548971

**Authors:** Yiheng Mao, Ying Chen, Yuan Li, Longda Ma, Xi Wang, Qi Wang, An He, Xi Liu, Tianyi Dong, Weina Gao, Yanfen Xu, Liang Liu, Liang Ren, Qian Liu, Peng Zhou, Ben Hu, Yiwu Zhou, Ruijun Tian, Zheng-Li Shi

## Abstract

The lung, as a primary target of SARS-CoV-2, exhibits heterogeneous microenvironment accompanied by various histopathological changes following virus infection. However, comprehensive insight into the protein basis of COVID-19-related pulmonary injury with spatial resolution is currently deficient. Here, we generated a region-resolved quantitative proteomic atlas of seven major pathological structures within the lungs of COVID-19 victims by integrating histological examination, laser microdissection, and ultrasensitive proteomic technologies. Over 10,000 proteins were quantified across 71 dissected FFPE post-mortem specimens. By comparison with control samples, we identified a spectrum of COVID-19-induced protein and pathway dysregulations in alveolar epithelium, bronchial epithelium, and pulmonary blood vessels, providing evidence for the proliferation of transitional-state pneumocytes. Additionally, we profiled the region-specific proteomes of hallmark COVID-19 pulmonary injuries, including bronchiole mucus plug, pulmonary fibrosis, airspace inflammation, and hyperplastic alveolar type 2 cells. Bioinformatic analysis revealed the enrichment of cell-type and functional markers in these regions (e.g. enriched TGFBI in fibrotic region). Furthermore, we identified the up-regulation of proteins associated with viral entry, host restriction, and inflammatory response in COVID-19 lungs, such as FURIN and HGF. Collectively, this study provides spatial proteomic insights for understanding COVID-19-caused pulmonary injury, and may serve as a valuable reference for improving therapeutic intervention for severe pneumonia.

## INTRODUCTION

The coronavirus disease 2019 (COVID-19) pandemic, caused by the severe acute respiratory syndrome coronavirus 2 (SARS-CoV-2), remains a severe threat to public health.^1^ It has led to over 767 million confirmed cases and 6.94 million deaths worldwide as of July 1, 2023. While the majority of COVID-19 patients were either asymptomatic or experienced mild symptoms, a considerable proportion of patients progressed to severe disease or even death.^2^ Despite the efforts to develop new vaccines and targeted drugs, the pandemic is still spreading without sufficient therapeutic options available for severe cases of COVID-19.^3^ One key challenge is the incomplete understanding of the mechanisms underlying COVID-19-associated multi-organ injuries and failures. Therefore, further investigation into the pathological and molecular basis of COVID-19 pneumonia is warranted to develop more effective treatments, particularly for critical cases.

Extensive organ injuries and systemic viral infection were observed in multiple human body systems of COVID-19 patients, among which the respiratory system is a primary target of SARS-CoV-2.^4^ Due to the vulnerability of human lungs, respiratory failure caused by acute lung injury or acute respiratory distress syndrome (ARDS) becomes a leading cause of COVID-19-related fatalities.^5^ The normal human lung has three essential structures including pulmonary alveoli, respiratory bronchioles, and blood vessels to maintain the gas exchange function. Following infection by SARS-CoV-2, these structures are gradually damaged accompanied by the presence of a range of lesions.^6^ The main pulmonary pathological alterations of COVID-19 were manifested as diffuse alveolar damage (DAD), interstitial fibrosis, and exudative inflammation in autopsy studies using traditional tissue pathology techniques, such as light microscopy and immunohistochemistry (IHC).^7–9^ Nonetheless, the molecular basis underlying lung pathogenesis after viral infection remains to be fully elucidated.

After the pandemic outbreak, single-cell RNA sequencing (scRNA-seq) and digital spatial profiling (DSP) technologies were exploited to depict the high-resolution transcriptomic atlas of lung pathology during COVID-19 progression, achieving remarkable depth and throughput.^10–12^ Instead of RNA, the cellular machinery and biological processes altered by virus infection are primarily controlled by proteins. Accordingly, several research groups have provided protein-level insights into the pathogenesis of COVID-19 through profiling the global proteome of COVID-19 patient autopsies.^13–15^ These studies predominantly focused on the overall proteome changes by analyzing bulk tissue samples, which inevitably compromised the spatial and cell-type information and potentially brought biased results. In order to determine the proteome alteration in different structures of COVID-19 pulmonary injury, a comprehensive characterization of the spatial proteomic landscape of lung pathology in COVID-19 post-mortem specimens is warranted.^16^

In this study, we combined histopathological examination, single-cell-accuracy laser microdissection (LMD), a <u>s</u>imple and integrated <u>s</u>pintip-based <u>pro</u>teomics technology (SISPROT),^17, 18^ and an ultra-sensitive liquid chromatography-mass spectrometry (LC-MS) platform to construct a deep and spatially resolved proteomic atlas of severe COVID-19-related pulmonary injury. By enhancing the overall analytical sensitivity, we quantified more than 10,000 proteins in small-size tissue slices (i.e. an area of 1 mm^2^ for each sample) dissected from the formalin fixed paraffin-embedded (FFPE) lung autopsy specimens. We collected and analyzed three key pulmonary structures namely alveolar epithelium (AE), bronchial epithelium (BE), and blood vessels (VE), from both COVID-19 and non-COVID-19 autopsies. Additionally, we investigated the proteomes of four other hallmark pathological alterations including bronchiole mucus plug (BMP), pulmonary fibrosis (PF), airspace inflammation (ASI) and hyperplastic alveolar type 2 (HAT2) in COVID-19 lungs. Our dataset unveiled a range of region-specific in protein and pathway dysregulations induced by COVID-19 and also linked the pulmonary pathological lesions to spatially resolved proteomes. Through bioinformatic analysis, we discovered the spatial enrichment of key cell-type and functional protein markers, some of which were further validated using IHC and immunofluorescence (IF). Overall, this study provides a unique perspective on the characteristics and mechanisms of pulmonary injury in severe COVID-19 patients, and can serve as a valuable resource for identifying potential therapeutic target for the disease.

## RESULTS

### Generation of a pathology-guided and in-depth spatial proteomic atlas of COVID-19 lungs

To comprehensively investigate the region-resolved proteome of severe COVID-19-related pulmonary injury with limited starting material, a project-specific workflow was developed to increase the analytical sensitivity (**Figure 1A**). Firstly, an integrated proteomic technology called SISPROT developed by us previously was adapted for rare FFPE sample processing.^17–19^ The lysis buffer formula, heating duration, and cutting area were optimized to increase protein identification from cross-linked and hematoxylin and eosin (H&E) stained tissue sections (**Table S1** and **S2**). Secondly, the laser parameters of the LMD microscopy were optimized to reduce tissue damage caused by laser ablation (**Figure S1**). Thirdly, a high-speed timsTOF Pro MS was equipped with a narrow-bore C18 column (i.e. 50 μm inner diameter) for nanogram-level peptide sample analysis.^20^ Finally, the state-of-art parallel accumulation–serial fragmentation combined with data-independent acquisition (diaPASEF) method was employed to increase reproducibility, quantitative accuracy, and proteome coverage.^21^

**Figure 1.**
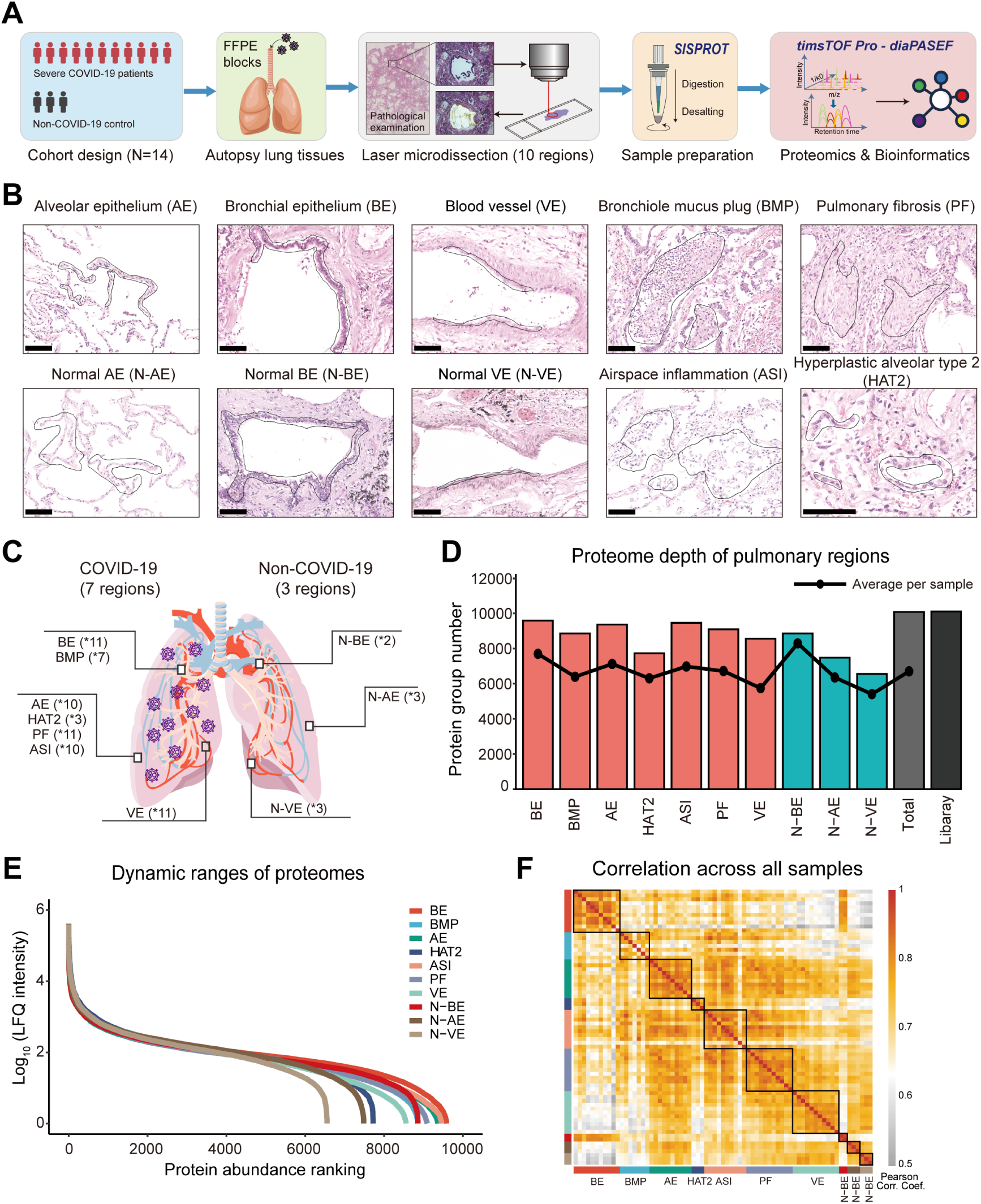
Spatial proteome profiling of pulmonary injury in COVID-19 autopsies. (A) Schematic experimental workflow including cohort design, autopsy specimen collection, pathology-guided laser microdissection, integrated proteomics sample preparation, mass spectrometry-based deep proteome profiling, and bioinformatic analysis to analyze the ten regions of human pulmonary tissues of severe COVID-19 patients and non-COVID-19 control individuals. (B) Representative hematoxylin and eosin (H&E) staining images of ten selected regions from COVID-19 and non-COVID-19 specimens for laser microdissection and following proteome profiling (scale bar = 50 μm). Region names and their abbreviations are labeled above images. (C) Schematic human pulmonary structure. Biological replicate counts for each region are labeled. (D) Total number (as bar graph) and average number (as black dot) of identified protein groups for each region and project-specific data-dependent acquisition-based library. (E) Dynamic range of the proteome within ten regions based on average protein label-free quantification (LFQ) intensity. (F) Heatmap of Pearson correlation coefficients of protein LFQ intensities across all samples.

Post-mortem lung samples in the form of FFPE blocks were obtained from 11 victims who succumbed to COVID-19 during the first-wave pandemic in Wuhan, China. The age of the patients was between 51 and 86 years. The patients were symptomatic for an average of ∼34.6 days (range, 17 to 89 days) prior to death (**Table S3**). To make a direct comparison between corresponding structures in COVID-19 and non-COVID-19 lungs, we also collected FFPE specimens of healthy lungs from three other victims of traffic accidents or coronary heart disease. Following a thorough pulmonary histopathology examination, we selected and dissected ten regions of interest (ROIs) (**Figure 1B**, **1C** and **S2**). These includes three basic pulmonary structures from COVID-19 specimens namely AE (n = 10), BE (n = 11), and VE (n = 11), as well as paired morphologically normal non-COVID-19 samples namely N-AE (n = 3), N-BE (n = 2), and N-VE (n = 3) (**Table S4**). Additionally, we also collected four crucial pathological alteration namely BMP (n = 7), PF (n = 11), ASI (n = 10), and HAT2 (n = 3) for subsequent spatial proteomic analysis.

This combinatory spatial proteomic workflow allowed us to map the expression of thousands of proteins in single-shot LC-MS analysis from 1 mm^2^ of FFPE tissue section in an unbiased fashion. Initially, a project-specific spectral library was generated by high-pH reversed-phase fractionation of pooled samples and repeated injection of LMD samples. The resulting hybrid library comprised 210,597 precursors, accounting for 168,887 peptides and 11,438 protein groups, which provided a comprehensive reference database for the following DIA search. To profile the spatial proteome, an accumulated area of 1 mm^2^ (5 μm in thickness) of each region was collected by LMD, followed by SISPROT processing and DIA measurement. In total, we quantified 10,075 protein groups from 71 samples of ten regions (**Figure 1D**). The proteome depth of our dataset is comparable to that of reported bulk tissue analysis using hundreds of micrograms of starting material *via* a conventional pipeline. More than 8,500 protein groups were quantified in average in ten pulmonary regions (i.e., 9,583 in BE, 8,854 in BMP, 9,364 in AS, 7,725 in HAT2, 9,460 in ASI, 9,091 in PF, 8,556 in VE, 8,858 in N-BE, 7,482 in N-AS, and 6,554 in N-VE). For 71 DIA runs, an average of 6,700 protein groups were quantified per sample (**Figure S3A**). The detailed “protein intensity in ROIs” information for each identified protein were provided in **Supplementary Data 1**. The label-free quantification (LFQ) intensity of proteins identified in ten pulmonary regions spanned near six orders of magnitude and showed a moderately high intra-region correlation (**Figure 1E** and **1F**). Notably, the intensity correlations of non-COVID-19 control samples were higher than COVID-19 ones (**Figure S3B**), which may be attributed to the heterogeneous nature of virus-infected organs, even in the same pathology-defined region.^22^ To ensure the reproducibility of sample preparation and stability of LC-MS system, we processed six replicates of 100 ng HEK 293T cell lysates along with LMD samples. The LFQ intensity of proteins identified in these technical replicates showed relatively low median coefficient of variation (CV) of 0.13 (**Figure S4**). Overall, these results demonstrated the effectiveness of our spatial proteomic workflow.

### Comparative analysis reveals dysregulation of proteins and related pathways in three basic pulmonary structures

Although global proteome changes of SARS-CoV-2-infected human lungs have previously been described through bulk tissue analysis elsewhere, we aimed to identify and locate dysregulated proteins that may have been averaged in other studies.^13–15^ To characterize the region-specific proteomic change and dysregulation of related functions and pathways caused by COVID-19 in post-mortem lungs, we conducted a comparative proteome analysis of the three basic pulmonary structures (i.e., AE, BE, and VE) between COVID-19 samples and non-COVID-19 controls. As expected, the COVID-19 samples can be well distinguished from non-COVID-19 controls in principle component analysis (PCA) for all three regions (**Figure 2A**), indicating distinct proteomic features of the two groups. Subsequently, we performed two-sample t-test to filter differently expressed proteins (p-value < 0.05, fold change of LFQ intensity > 2). There were 397, 487, and 437 significantly up-regulated proteins, and 1,089, 345, and 475 significantly down-regulated proteins identified in the AE, BE, and VE regions of COVID-19 samples compared with the non-COVID-19 controls, respectively (**Figure 2B** and **Table S5**).

**Figure 2.**
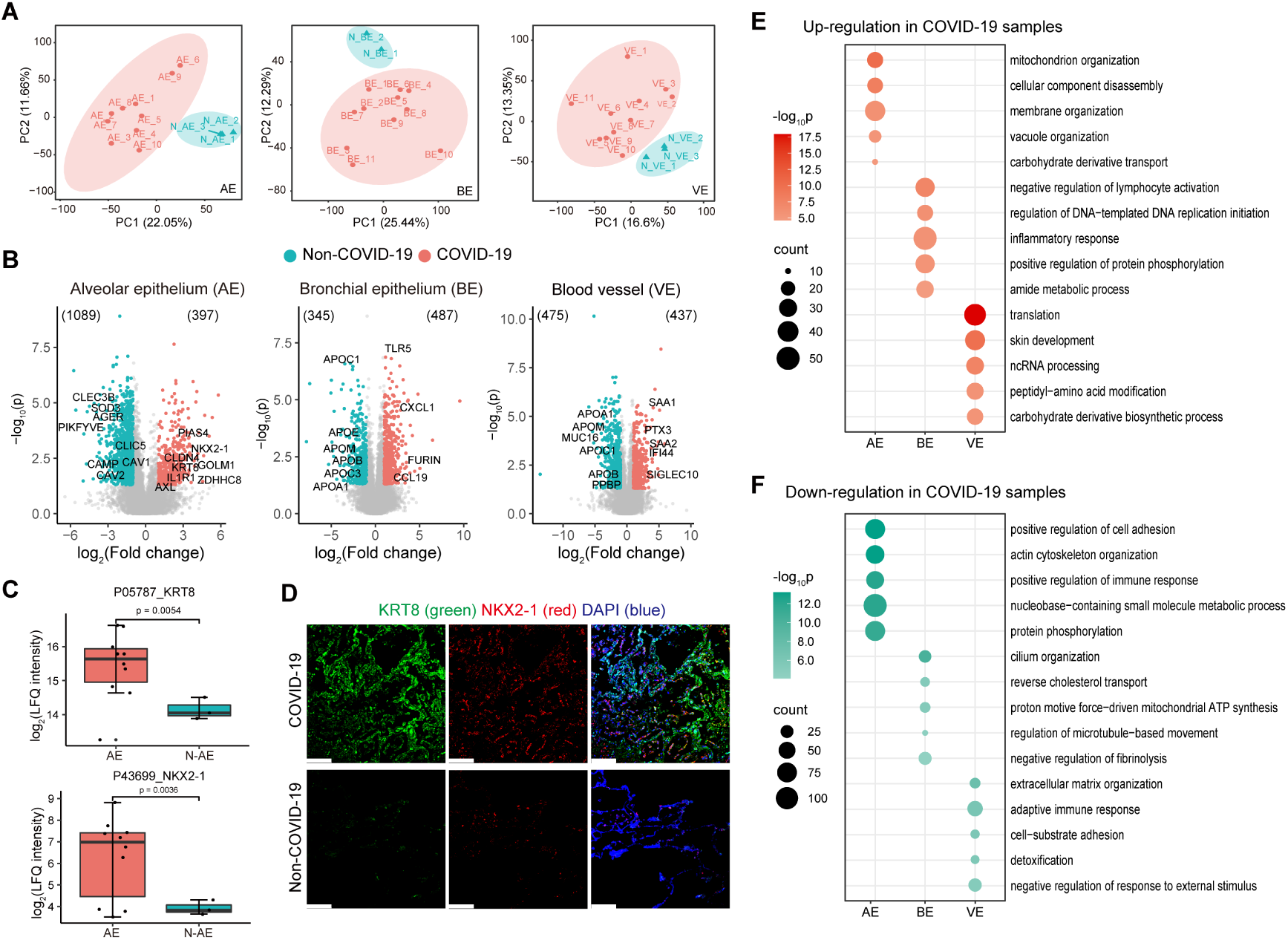
Region-specific dysregulation of protein expression in three basic pulmonary structures of COVID-19 lungs. (A) Principal component analysis (PCA) of the proteome profiles in the three basic structures of lungs from with severe COVID-19 patients and non-COVID-19 control individuals. (B) Volcano plots of the COVID-19 samples versus non-COVID-19 samples are shown for the three basic pulmonary structures including alveolar epithelium (AE), bronchial epithelium (BE), and blood vessels (VE). Differentially expressed proteins (p-value < 0.05 and LFQ intensity of COVID-19 / non-COVID-19 > 2 or < 1/2) are highlighted in pink (up-regulated in COVID-19 group) and cyan (down-regulated in COVID-19 group). (C) Box-whisker plots of protein expression of KRT8 and NKX2-1 in AE regions from COVID-19 lungs and non-COVID-19 lungs. (D) Immunofluorescence images of KRT8 and NKX2-1 in AE regions from COVID-19 lungs and non-COVID-19 lungs (scale bar = 50 μm). (E) Dot plot showing the top-5 GOBP terms enriched in significantly up-regulated proteins of AE, BE, and VE regions in COVID-19 lungs. (F) Dot plot showing the top-5 GOBP terms enriched in significantly down-regulated proteins of AE, BE, and VE regions in COVID-19 lungs.

AE constitute as a thin wall responsible for gas exchange in the lung. Alveolar damage has been extensively observed in COVID-19 autopsy studies with evidence shown that SARS-CoV-2 predominantly infects alveolar epithelial cells and leads to AE cell number reduction.^22, 23^ Recent studies have identified a transitional population of alveolar epithelial cells, marked by elevated expression of KRT8, CLDN4, and IL1R1, which dramatically increased in response to pulmonary injury in animal models. Interestingly, we also detected the up-regulation of these three protein markers in the COVID-19 AE (**Figure 2B**), confirming the existence and expansion of so-called “pre-alveolar type 1 transitional cell state (PATS)” in patients’ lungs affected by COVID-19.^24–27^ Besides, the PATS cells were postulated to be cancer cell-like due to their accumulation in cell number and DNA damage in previous report.^24^ Surprisingly, our data also revealed an increase of NKX2-1 (also known as TTF1), an important oncogene in lung adenocarcinomas, in COVID-19 AE, providing further support for this hypothesis.^28^ Furthermore, we validated our proteomic discovery by IF staining (**Figure 2C** and **2D**) which confirmed the increased expression of KRT8 and NKX2-1 in COVID-19 AE region. In addition, we also observed the up-regulation of several potential drug targets of SARS-CoV-2 in the AE region, including ZDHHC8 (a palmitoyltransferase critical for spike protein-mediated virus entry^29^), PIAS4 (a SUMO ligase capable of stabilizing ACE2^30^), GOLM1 (a glucogenic hormone contributing to SARS-CoV-2-induced hyperglycemia^31^), and AXL (a candidate receptor for SARS-CoV-2^32^) in COVID-19 AE. As for the down-regulated proteins in AE, we detected PIKFYVE a well-known drug target for viral entry and replication^33^, as well as the antioxidant enzyme SOD3, the tetranectin CLEC3B, and the antimicrobial peptide CAMP, which were reported to decrease in COVID-19 lung tissue or plasma.^34–36^ Furthermore, we observed a decrease in four human lung alveolar type 1 (AT1) cell markers, AGER, CAV1, CAV2, and CLIC5, in the AE, suggesting a loss of normal alveolar function in COVID-19.^37^

BE is comprised of basal, ciliated, and secretory cells and serves as a defensive barrier in the airway, making it a major target of inhaled pathogen. In the BE of COVID-19 lungs, we observed the up-regulation of important chemokines such as C-X-C motif chemokine ligand 1 (CXCL1), and C-C motif chemokine 19 (CCL19) which is in line with previous transcriptomic studies^38, 39^, signifying heightened inflammatory tissue damage, lung injury, and respiratory failure (**Figure 2B**). Our findings suggest that inhibition of these pathways could suppress immune hyperactivation in severe COVID-19 cases. Moreover, we identified the up-regulation of Toll-like receptor 5 (TLR5), a key component of the innate immune response, as well as the protease FURIN, which is implicated in SARS-CoV-2 invasion. These proteins can be considered as potential therapeutic targets for future drug development.^40, 41^ For the down-regulated proteins, we identified several apolipoproteins involved in lipid metabolism, including APOA1, APOB, APOC1, APOC3, APOE, and APOM. Similar reductions in apolipoproteins have been identified in the serum of severe COVID-19 patients.^42^ Notably, APOE plays a role in mediating the inflammatory response in BE, and has been confirmed to interact with ACE2, thereby inhibiting SARS-CoV-2 cellular entry and inflammation in COVID-19 patients.^43, 44^ Our data further strengthens the connection between apolipoproteins and the severity of COVID-19.

VE plays a key role in the circulation of blood and encompasses a number of cell types. The VE region dissected in this study comprised vascular endothelium and surrounding elastic walls. Histopathological analysis of pulmonary vessels in COVID-19 autopsies revealed extensive thrombosis, endothelialitis, and angiogenesis.^45^ The injury to VE can sometimes lead to life-threatening complications^46^; however, the mechanisms underlying this dysfunction have not been fully elucidated. We observed the up-regulation of activated acute phase proteins SAA1 and SAA2 in COVID-19 VE (**Figure 2B**). These proteins have also been found to be enriched in the serum of severe COVID-19 patients and show potential as biomarkers for viral infection.^42^ Additionally, we observed the up-regulation of several proteins that are known to be associated with SARS-CoV-2 invasion, including SIGLEC10 (a receptor binding domain (RBD) interacting protein^47^), IFI44 (an immune evasion biomarker for SARS-CoV-2^48^), and PTX3 (a predictor of COVID-19 mortality^49^). We also identified down-regulated proteins within the VE, namely MUC16, PPBP, APOM, APOA1, APOC1, and APOB, which is consistent with findings from previous serum transcriptomics and proteomic studies.^42, 50^ The results above suggested a disordered inflammation in COVID-19 VE.

After analyzing differentially expressed proteins individually, we performed the gene ontology biological processes (GOBP) enrichment analysis of the six region-unique dysregulated proteomes using the Metascape platform^51^ (**Supplementary Data 2**). The top-5 significantly up-regulated and down-regulated biological processes in AE, BE, and VE were shown in **Figure 2E** and **2F**, respectively. We observed distinct and shared dysfunctions caused by COVID-19 across the three pulmonary structures. For example, we found significant up-regulation of cellular components disassembly and mitochondrion organization in AE, and down-regulation of cilium organization in BE. These findings suggest that SARS-CoV-2 can result in widespread tissue damage and loss of normal pulmonary functions. More importantly, a series of immune response-related biological processes were found to be significantly enriched. In general, the lymphocyte activation and adaptive immune response were negatively regulated in COVID-19 lungs, while the inflammatory response was up-regulated. This aligns with clinical reports of lymphopenia, impaired adaptive immune responses, and disordered expression of cytokines and chemokines in severe COVID-19 patients^52, 53^, further emphasizing the importance of targeted therapy against immune system disorders that cause multi-organ failures in fatal cases of COVID-19.

### Spatial proteome and functional characteristics of seven COVID-19 pulmonary regions

Following the comparative analysis of three basic pulmonary structures, we proceeded to investigate the proteomic characteristics of four other hallmark pathological regions of COVID-19 lungs including BMP, PF, ASI, and HAT2. Due to the deficiency of corresponding regions in healthy lungs, it was a challenge to obtain enough material as control group for these four regions. Consequently, here we focus on analyzing COVID-19 samples to explore their cell-type and functional differences. By conducting analysis of variance (ANOVA) and Gene Ontology Biological Process (GOBP) enrichment, we identified seven protein clusters characterizing the different regions of COVID-19 lungs with distinct functions (**Figure 3A** and **Table S6**), including cluster 1 for BE (1,678 proteins), cluster 2 for BMP (659 proteins), cluster 3 for AE (139 proteins), cluster 4 for HAT2 (749 proteins), cluster 5 for ASI (22 proteins), cluster 6 for PF (148 proteins), and cluster 7 for VE (800 proteins).

**Figure 3.**
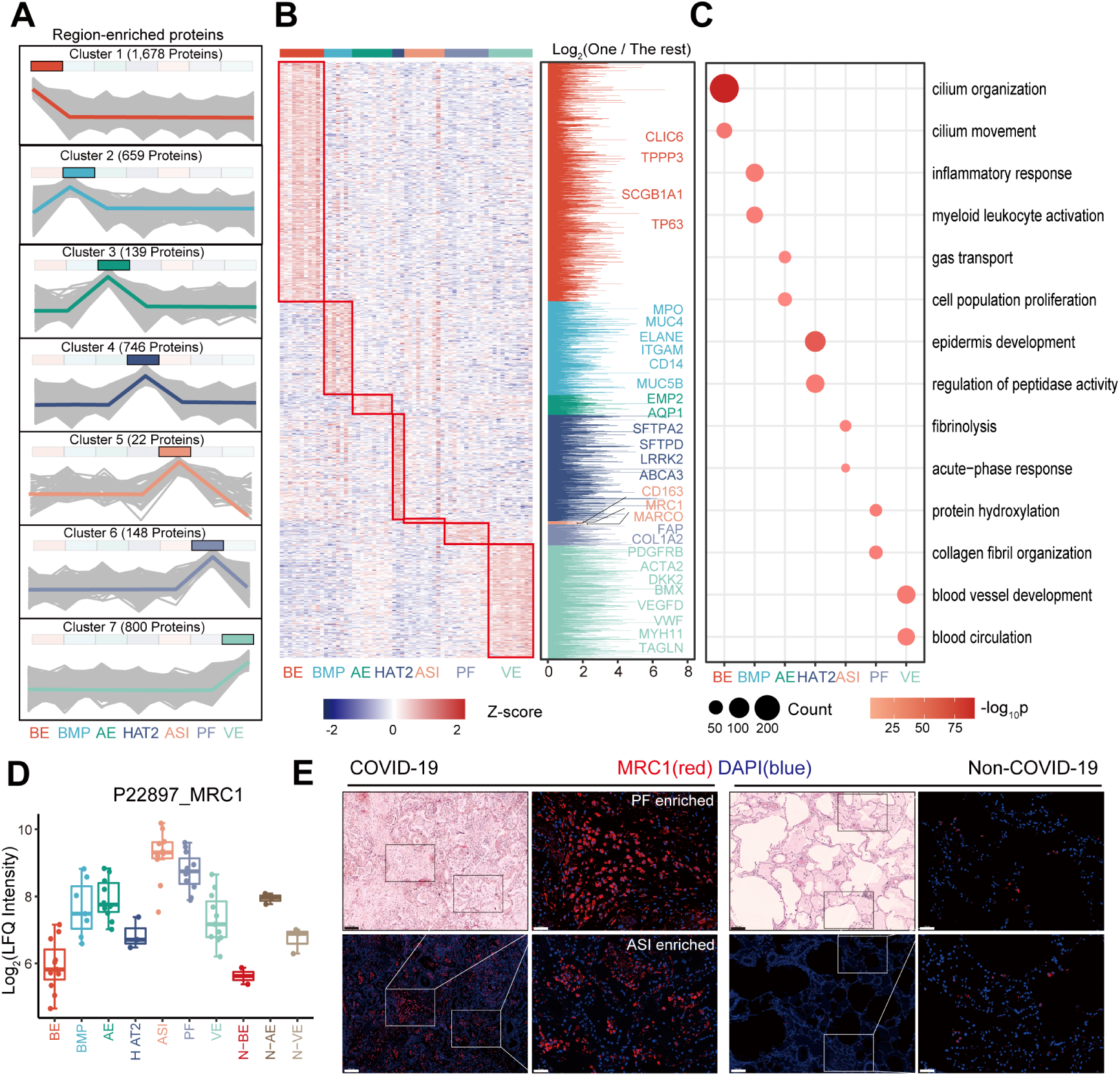
Proteomic and functional features of hallmark lesions in COVID-19 lung autopsies. (A) Seven protein expression clusters enriched in seven pulmonary regions by co-expression analysis, including BE (Cluster 1), BMP (Cluster 2), AE (Cluster 3), HAT2 (Cluster 4), PF (Cluster 5), ASI (Cluster 6), and VE (Cluster 7). Log_2_ fold change (FC) was calculated by the median of proteins expression ratio between one region and the rest. Proteins with FC larger than 1.2 were kept for following analysis. The proteins with the one-way analysis of variance (ANOVA) p-value < 0.05, FDR < 0.05 were regarded as statistically significant in the specific region. (B) Co-expression patterns for region-enriched proteins in the seven clusters, respectively. Left panel: heatmap showing the Z-score normalized protein LFQ intensities of region-enriched proteins in each sample. Right panel: histogram represents up-regulated proteins in the seven regions. The protein names of representative cell-type and functional markers of each region are labeled. (C) Dot plot showing selected GOBP terms for each region-enriched protein cluster. The size of circle represents the number of proteins and the color indicates the -log_10_ p-value. (D) Box-whisker plot showing protein expression of MRC1 in the ten regions from COVID-19 and non-COVID-19 lungs. (E) Immunofluorescence images of MRC1 in COVID-19 lungs and non-COVID-19 lungs with top left panels showing H&E stained images of the same region in an adjacent tissue section (scale bar = 50 μm).

Initially, we checked if defined cell type markers can be identified in respective clusters (**Figure 3B**).^11, 12, 37^ As expected, airway epithelial marker TP63, CLIC6, TPPP3, and SCGB1A1 were enriched in Cluster 1 (BE); myeloid cell marker MPO, ELANE, CD14, and ITGAM, and mucin MUC5B and MUC4 were enriched in Cluster 2 (BMP); AT1 cell marker EMP2 and AQP1 were enriched in Cluster 3 (AE); alveolar type 2 (AT2) cell marker SFTPA2, SFTPD, ABCA3, and LRRK2 were enriched in Cluster 4 (HAT2); macrophage marker CD163, MARCO, and MRC1 were enriched in Cluster 5 (ASI); fibroblast marker FAP, FN1, LUM, MMP2, and MMP14 and collagen deposition marker COL1A2, COL3A1, COL5A1, and COL12A1 were enriched in Cluster 6 (PF); vascular endothelial marker VEGFD, VWF, BMX, and DKK2, as well as vascular surrounding stroma marker PDGFRB, ACTA2, TAGLN, and MYH11 were enriched in Cluster 7 (VE). Furthermore, GOBP analysis revealed enriched processes specific to each region (**Figure 3C**). For instance, Cluster 1 (BE) was involved in cilium organization and cilium movement; Cluster 3 (AE) was involved in gas transport and cell population proliferation; and Cluster 7 (VE) was involved in blood vessel development and blood circulation. Taken together, these results uncovered distinct proteomic characteristics and functional differences among the seven COVID-19 pulmonary regions and well demonstrated the reliability of our spatial proteomic dataset.

Notably, Cluster 5 (ASI) has the lowest protein number among the seven clusters, indicating its relatively lower uniqueness in terms of protein components (**Figure 3A**). ASI is a hallmark of DAD in COVID-19 pneumonia, which can be reflected as ground-glass opacity by chest computed tomography (CT) on admission.^54^ In this region, we identified 22 enriched proteins (**Table S6**), including thrombosis-related proteins (i.e. F2, FGA, FGB, FGD2, FGG, PLG), lipid homeostasis-related proteins (i.e. CYP27A1, NCEH1, PLA2G15), and inflammatory-related proteins (i.e. IFI30, LILRB5, VSIG4). According to GOBP and Kyoto Encyclopedia of Genes and Genomes (KEGG) annotation, the ASI-enriched proteins were mainly involved in fibrinolysis, acute-phase response, lipid catabolic process, innate immune response, and phagosome (**Figure S5**). In addition, we identified three markers of proliferating alveolar macrophage in this cluster, including CD163, MARCO, and MRC1, confirming the infiltration of macrophages in damaged alveoli of COVID-19 lungs.^55, 56^ We validated the spatial distribution of mannose receptor MRC1 using IF staining. As shown in **Figure 3E**, MRC1 is significantly enriched in the ASI and PF regions of COVID-19 lung tissue, while showing minimal signal in the non-COVID-19 control sample. Monocyte-derived macrophages typically reside in alveoli and play an important role in COVID-19 pathogenesis.^57, 58^ Previous studies have reported that SARS-CoV-2 infection triggers the accumulation and activation of macrophages, driving profound pulmonary fibrosis in COVID-19 lungs. Our data provide further evidence for the enrichment of not only profibrotic macrophages but also related potential therapeutic target such as VSIG4 in COVID-19 autopsy lungs.^59^

### Proteomic and functional features of PF, BMP, and HAT2 in COVID-19 lungs

Fibrosis is a common occurrence in a spectrum of lung diseases, such as idiopathic pulmonary fibrosis (IPF) and lung cancer, which can lead to pulmonary architecture damage, progressive respiratory failure, and lethal hypoxemia.^60, 61^ In addition to severe cases of COVID-19, pulmonary fibrosis is often diagnosed as a critical sequela in COVID-19 survivors and long COVID-19 patients.^62, 63^ One main reason for fibrosis in COVID-19 lungs is the failure of pulmonary injury repair, which is triggered by persistent and reciprocal circuits between macrophages and fibroblasts, along with dysregulated signaling pathways, such as innate immune cell activation.^56^ Previous studies have shown that the transcriptional profile of COVID-19-induced pulmonary fibrosis is similar to that of IPF, which is short of effective therapeutic strategy currently.^55, 64^ Therefore, gaining a deeper understanding of the molecular mechanisms and etiology of COVID-19-induced pulmonary fibrosis is necessary to identify potential drug targets. In our dataset, 148 proteins were found to be enriched in the cluster of this region (**Figure 3A**), which were then subjected to GOBP and KEGG analysis. As expected, a range of biological processes and signaling pathways associated with excessive collagen deposition and extracellular matrix (ECM) production were significantly enriched, such as protein hydroxylation, collagen fibril organization, collagen metabolic process, cell-substrate adhesion, glycoprotein metabolic process, regulation of extracellular matrix disassembly, and tissue remodeling (**Figure 4A** and **4B**). Notably, we identified CTHRC1 in the “tissue remodeling” term, which was recently described as a hallmark gene of pathological fibroblasts contributing to rapid fibrosis in both IPF and COVID-19^11, 65^ (**Table S6** and **Supplementary Data 3**).

**Figure 4.**
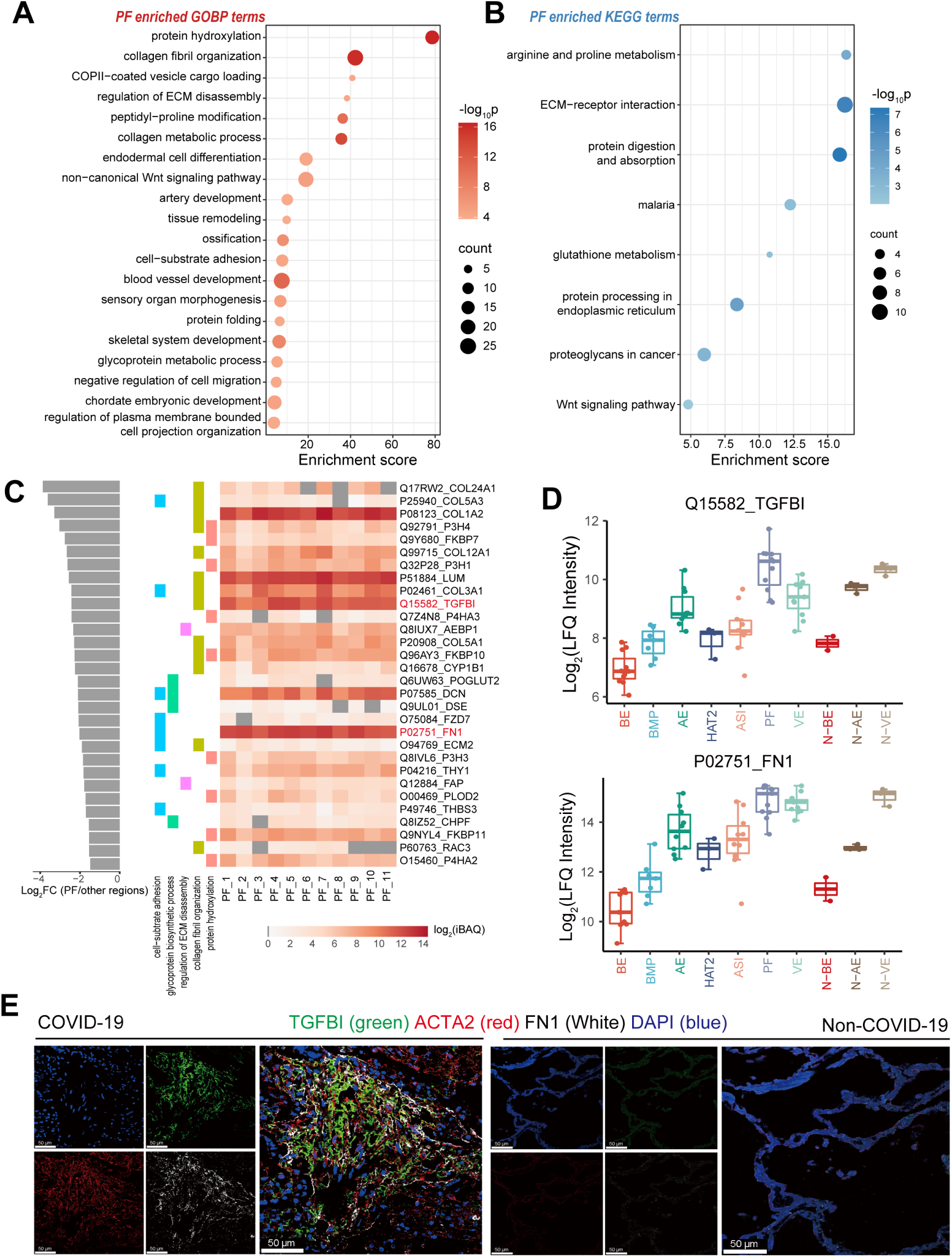
Proteomics-based functional analysis of pulmonary fibrosis in COVID-19 lungs. (A) Dot plot showing top-20 significant GOBP terms for PF-enriched protein cluster. The size of circle represents the number of proteins and the color indicates the -log_10_ p-value. (B) Dot plot showing all significant KEGG terms for PF-enriched protein cluster. The size of circle represents the number of proteins and the color indicates the -log_10_ p-value. (C) Protein expression patterns of selected GOBP terms. Left panel: log_2_FC was calculated by the median of proteins expression ratio between one and the rest regions. Proteins with top30 log_2_FC value are shown. Middle panel: annotated GOBP terms of each protein. Right panel: heatmap showing protein intensities of the 30 proteins in 11 PF biological replicates. (D) Box-whisker plots of protein expression of TGFBI and FN1 in the ten regions from COVID-19 and non-COVID-19 lungs. (E) Immunofluorescence images of TGFBI and FN1 in COVID-19 PF region and non-COVID-19 alveolar region with ACTA2 also being stained to indicate the location of PF in COVID-19 lungs (scale bar = 50 μm).

In order to identify potential vital protein regulators and markers in PF, we selected five representative GOBP terms and ranked involved proteins by their intensity fold changes (PF vs other regions). Among the top 30 proteins (**Figure 4C**), we found the collagens COL1A2, COL12A1, COL3A1, and COL5A1, collagen hydroxylases P3H4, P3H1, P3H3, and P4HA2, peptidyl-prolyl isomerases FKBP7, FKBP10, and FKBP11, along with some known ECM components and regulators including LUM, AEBP1, DCN, ECM2, THY1, FAP, PLOD2, and THBS3 in all seven biological replicates. Part of these proteins have the potential to serve as potential anti-fibrosis targets for future drug development. Intriguingly, we identified the transforming growth factor-beta-induced protein ig-h3 (TGFBI) and epithelial–mesenchymal transition (EMT) marker fibronectin FN1 with high protein intensities in this list (**Figure 4D**). TGFBI is an ECM glycoprotein which has been reported to be implicated in a variety of pathogenesis and biological processes^66, 67^, but its association with COVID-19 has not yet been explored in the literature. To validate our proteome profiles, we performed co-staining for FN1, TGFBI, and the myofibroblast marker ACTA2 (to locate the PF region) on tissue sections. As shown in **Figure 4E**, all these three proteins enriched and colocalized in the PF region in COVID-19 lung, while showing minimal signal in the alveolar region of control sample. These findings suggest that TGF-β pathway and EMT process play important roles in COVID-19 induced PF, highlighting the potential for targeted interventions to prevent and reverse PF.^60^

The normal function of airway mucus is to trap and expel inhaled pathogens and toxins for respiratory system defense.^68^ Excessive mucus accumulation and impaired clearance can lead to abnormal lung function, such as silent hypoxia and a decrease in pulmonary capacity in individuals with COVID-19.^69^ Previous studies have identified mucin and infiltrated innate immune cells as the main dysregulated components of BMP in COVID-19 lungs.^70, 71^ Moreover, proteomic changes of airway mucus isolated from COVID-19 patients by bronchoscopy have been investigated.^72^ In this study, we characterized BMP in COVID-19 lungs using LMD for better spatial accuracy. The cluster of BMP enriched 659 proteins against other regions (**Figure 3A**). We performed GOBP and KEGG analysis to determine function enrichment in BMP. Consistent with previous findings, a series of immune response-related biological processes were significantly enriched, such as leukocyte activation, regulation of defense response, immune effector process, innate immune response, and positive regulation of reactive oxygen species (ROS) (**Figure 5A** and **5B**). More importantly, multiple cytokine storm-related signaling pathways were observed including chemokine signaling pathway, natural killer cell mediated cytotoxicity, and IL-17 signaling pathway. These results provide valuable insights for the development of immunomodulatory treatments targeting the imbalanced inflammatory response specific to COVID-19.^73^

**Figure 5.**
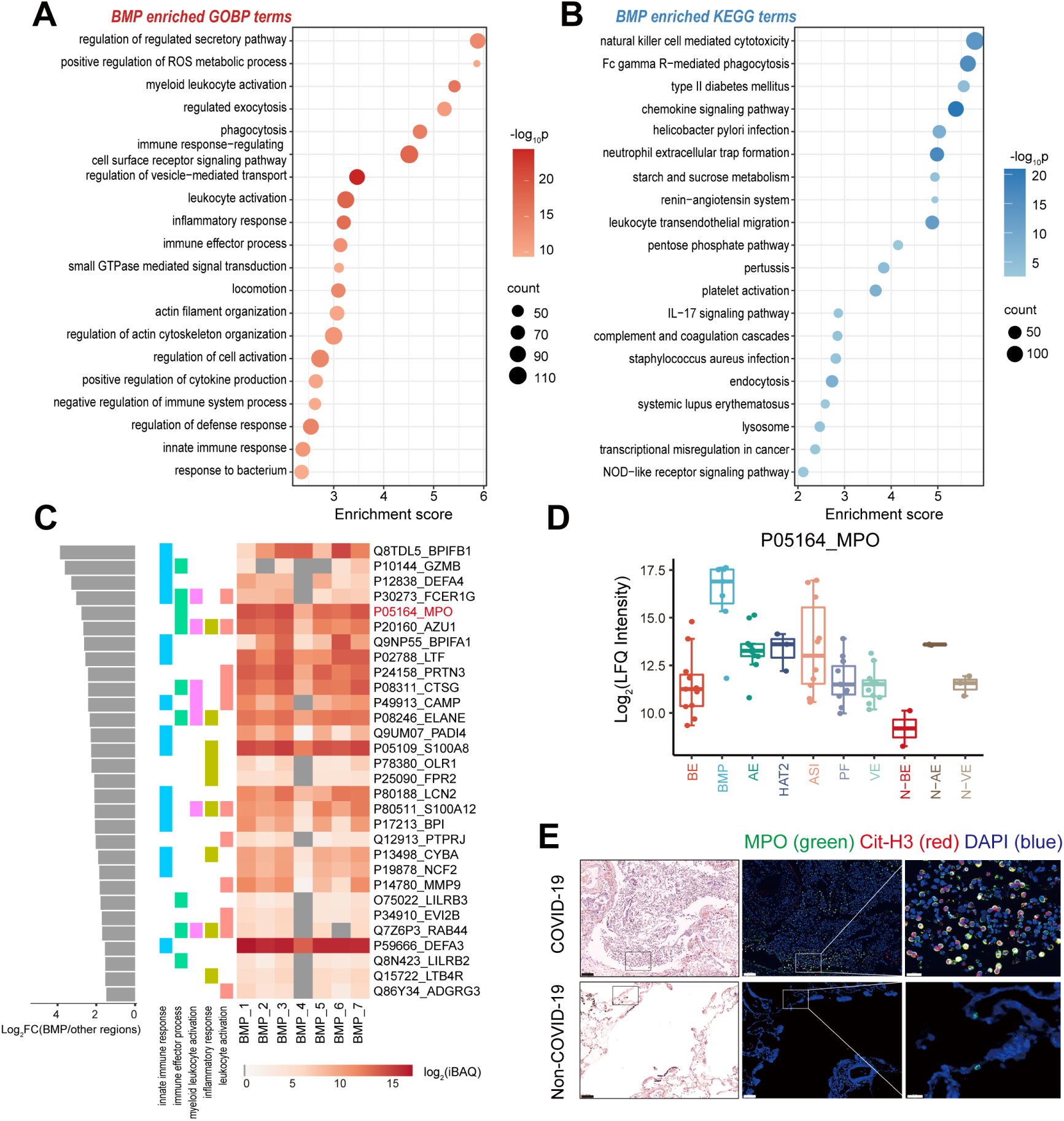
Proteomics-based functional analysis of bronchiole mucus plugs in COVID-19 lungs. (A) Dot plot showing top-20 significant GOBP terms for BMP-enriched protein cluster. The size of circle represents the number of proteins and the color indicates the -log_10_ p-value. (B) Dot plot showing top-20 significant KEGG terms for BMP-enriched protein cluster. The size of circle represents the number of proteins and the color indicates the -log_10_ p-value. (C) Protein expression patterns of selected GOBP terms. Left panel: log_2_FC was calculated by the median of proteins expression ratio between one and the rest regions. Proteins with top30 log_2_FC value are shown. Middle panel: annotated GOBP terms of each protein. Right panel: heatmap showing protein intensities of the 30 proteins in 7 BMP biological replicates. (D) Box-whisker plots of protein expression of MPO in the ten regions from COVID-19 and non-COVID-19 lungs. (E) Immunofluorescence images of MPO and Cit-H3 in COVID-19 BMP region and non-COVID-19 bronchiole region (scale bar = 50 μm).

The dysregulation of mucins in BMP has been well investigated in previous studies^69, 74^, so we focused on immune response-related proteins in this study. We selected five representative GOBP terms and ranked proteins involved by their intensity fold changes. In the top-30 list (**Figure 5C**), we identified several neutrophil markers including MPO, ELANE, DEFA3, and NCF2, as well as multiple functional proteins associated with neutrophils, including AZU1, PRTN3, CTSG, PADI4, S100A8, S100A12, and LILRB2 in all seven biological replicates, indicating the existence of high abundance of neutrophils in BMP. In addition, we observed a significant enrichment of the signaling pathway related to neutrophil extracellular trap (NET) formation (**Figure 5B**). The formation and release of NETs by neutrophils in response to viral infection has been found to be dysregulated in the blood and lungs of COVID-19 patients, potentially contributing to hyperinflammation and thrombosis.^75, 76^ To validate our proteome profiles, we co-stained for MPO and NET formation marker citrullinated histone H3 (Cit-H3) on tissue sections. As shown in **Figure 5E**, both markers were highly enriched and co-localized in the BMP region within the COVID-19 lung samples, while showing minimal signal in the bronchial region of the control sample. These results indicate the significant involvement of neutrophils and the NET process in the formation of the BMP and the dysregulated inflammatory response observed in COVID-19. Consequently, the inhibition of NET formation may provide potential relief for severe symptoms of COVID-19.

In healthy human lungs, AT2 cells primarily contribute to alveolar epithelial regeneration and surfactant secretion.^77^ When pulmonary injury occurs, AT2 can undergo proliferation and differentiate into AT1 cells as progenitor. However, this process can be hindered by SARS-CoV-2 infection. Histopathological examination studies of COVID-19 pulmonary tissues have revealed widespread hyperplasia of AT2 cells, which has been described as a hallmark of DAD.^8, 78^ Furthermore, AT2 cells tend to desquamate and cluster in COVID-19 lungs which made it possible for us to dissect and collected enough starting materials for proteome profiling.^23^ We performed GOBP analysis of the 749 proteins in HAT2 cluster and found the most significantly enriched process was epidermis development (**Figure S6**), suggesting that dysfunction of AT2 cells may result in keratinization. In addition to the aforementioned AT2 markers, we identified two reported virus entry-related protein including cathepsin CTSL and host cell receptor NPC1 enriched in HAT2, highlighting them as potential targets for anti-viral drug. Apart from CTSL, we also identified five other cathepsins enriched in HAT2, namely CTSD, CTSH, CTSV, and CTSS, which were involved in lysosome and apoptosis related pathways in KEGG analysis (**Table S6** and **Supplementary Data 3**). These findings may provide further insights into the mechanism of virus-induced HAT2 lesion.

### Multi-regional dysregulation of proteins related to viral entry and inflammatory response

In the analysis above, we profiled the region-specific proteomes of different pulmonary structures and described their enriched functions. However, there is a drawback that certain key proteins dysregulated in multiple regions may have been neglected. On the other hand, viral infection and activated immune response have been vastly observed in many cell types of COVID-19 lungs.^9^ Therefore, we focused on the dysregulation of proteins related to viral entry, host restriction, and inflammatory response across multiple pulmonary regions here. To facilitate our investigation, we constructed a manually curated database comprising proteins associated with COVID-19 pathogenesis, including virus receptors, viral entry cofactors, viral-entry-related proteases, and proteins involved in inflammation, based on reported and predicted datasets.^13, 79–83^ Out of the 273 proteins in this database, we identified 128 proteins in our data, with 34 proteins significantly up-regulated in the COVID-19 regions (**Figure 6A**; **Table S7** and **S8**).

**Figure 6.**
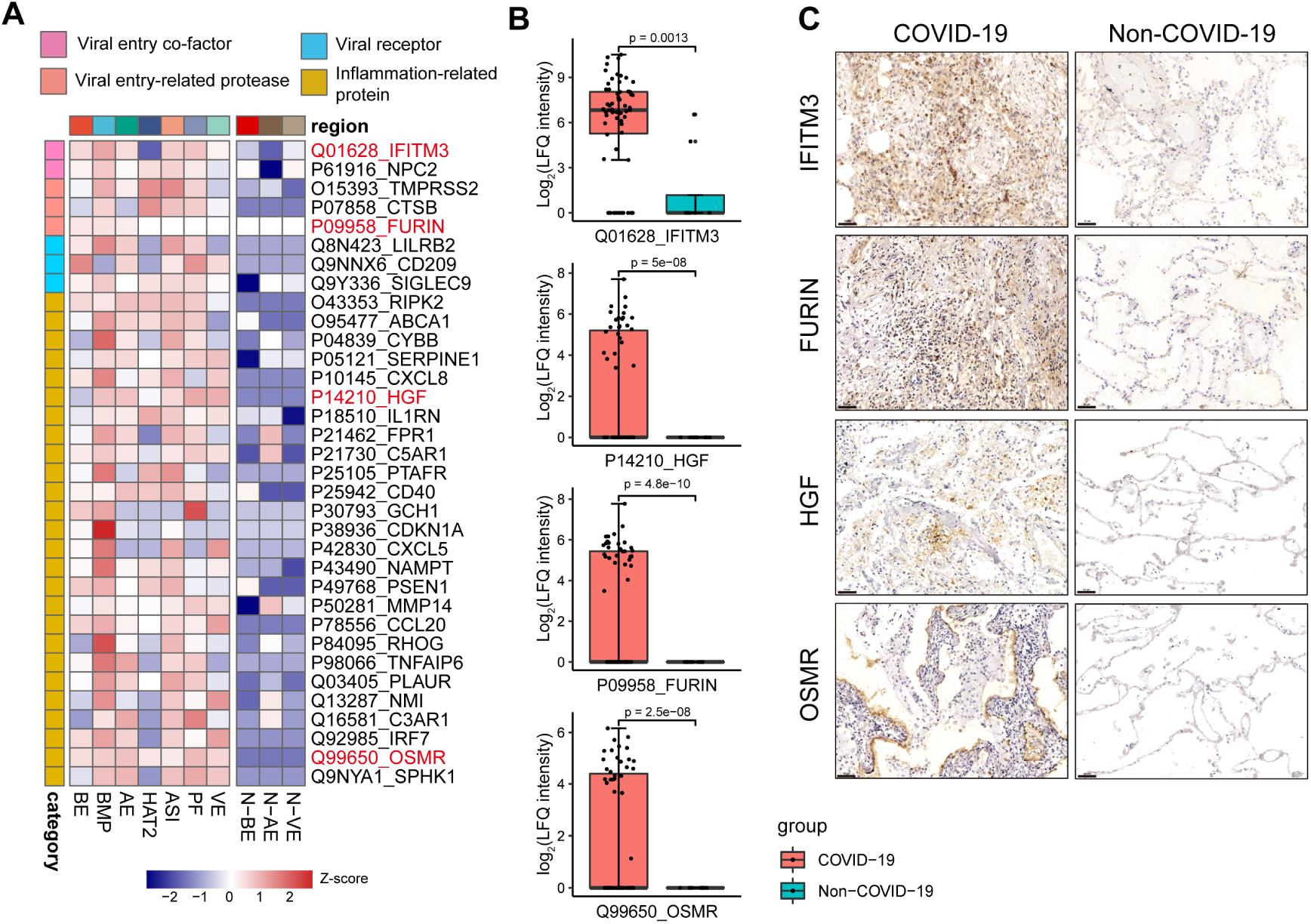
Dysregulated viral entry and inflammatory-related protein in COVID-19 lungs. (A) Heatmap showing the Z-score normalized intensities of significant up-regulated proteins involved in viral entry, host restriction, and inflammatory response in the ten regions of COVID-19 and non-COVID-19 lungs. (LIMMA, p-value < 0.05, Fold Change > 2) (B) Box-whisker plots of protein expression of IFITM3, HGF, FURIN, and OSMR in all samples from COVID-19 and non-COVID-19 lungs. (C) Immunohistochemistry images of IFITM3, HGF, FUSIN, and OSMR in COVID-19 lungs and non-COVID-19 lungs (scale bar: 50 μm).

Our data revealed the up-regulation of several viral infection-related proteins in COVID-19 lungs, including IFITM3, TMPRSS2, CTSB, FURIN, NPC2, LILRB2, CD209, and SIGLEC9. In the viral entry process, the interferon-induced transmembrane protein IFITM3 can interact and be hijacked by the SARS-CoV-2 spike (S) protein for efficient viral infection.^84^ In addition, proteases TMPRSS2 and FURIN can be implicated in the activation and cleavage of the S protein. While these findings were primarily derived from cell line or animal model experiments, our data provided further evidence for the critical roles of these proteins by proteomic profiling of autopsy specimens. Regarding proteins associated with inflammatory responses, we identified the up-regulation of chemokines CXCL8 and CXCL5, cytokines CCL20 and HGF, protein receptors FPR1, C3AR1, C5AR1, PTAFR, CD40, and PLAUR, as well as several key immunological regulators such as SERPINE1, IL1RN, and IRF7. Thereinto, C5AR1 and SERPINE1 have been suggested as potential therapeutic targets for alleviating excessive inflammation in severe COVID-19.^85, 86^ The expression levels of HGF, CCL20, and IRF7 have been reported to be associated with COVID-19 severity and mortality.^38, 87, 88^ Our data underscore the significance of these proteins in COVID-19 pathogenesis. Interestingly, we also observed the up-regulation of oncostatin-M-specific receptor OMSR, which involved in perturbed inflammation and fibroblast reprogramming in tumor growth^89, 90^, with few reports linking it to COVID-19 thus far. To valid our findings, we stained for IFITM3, FURIN, HGF, and OSMR on COVID-19 and non-COVID-19 pulmonary tissues. The IHC results are consistent with our proteome profiles (**Figure 6B** and **6C**).

## DISCUSSION

In the present study, we applied a highly sensitive MS-based spatially resolved tissue proteomics workflow for analyzing post-mortem specimens from COVID-19 victims. To the best of our knowledge, this study resulted in the deepest spatial proteome of COVID-19 samples to date, with more than 10,000 protein groups quantified in total. In addition, our workflow, especially the sample preparation technology SISPROT is scalable and user-friendly, allowing deep proteome profiling with limited starting materials. This capability could be beneficial for other proteomics studies at a nanogram level. Despite the depth of our data, we have not yet identified some known SARS-CoV-2 proteins and host receptors such as S protein and ACE2, which have also shown low expression levels in other COVID-19 tissue proteome resources.^13, 14^ The result may be attributed to decreased pulmonary viral load in the lungs of patients with a longer duration of the disease.^91^ Therefore, targeted assay, such as antibody-based staining or parallel reaction monitoring (PRM), can serve as a valuable complement for detecting these low-abundance proteins.

LMD-based proteomics has been successfully applied to unbiasedly analyze protein expression in tissue sections with spatial resolution in COVID-19 human hearts^16^, as well as other pulmonary diseases, such as tuberculosis granulomas and IPF.^92, 93^ In this study, we employed LMD technology to dissect and profile the proteomes of severe pulmonary injuries related to COVID-19 for the first time. Seventy-one samples of ten regions were collected, including three basic pulmonary structures from both COVID-19 and non-COVID-19 specimens, as well as four hallmark COVID-19 pulmonary pathological alterations. Comparative proteomic analysis of AE, BE, and VE between COVID-19 samples and non-COVID-19 controls revealed a series of COVID-19 induced protein and pathway dysregulations. For example, we detected the up-regulation of transitional-state pneumocyte markers (i.e. KRT8, CLDN4, and IL1R1) and down-regulation of normal human lung AT1 cell markers (i.e. AGER, CAV1, CAV2, and CLIC5) in COVID-19 AE, suggesting the excessive proliferation of transitional-state pneumocyte and loss of normal alveolar function in COVID-19. In BE and VE, dysregulation of immune response-related proteins and apolipoproteins was observed, indicating heightened inflammatory tissue damage, lung injury, and respiratory failure. Subsequently, we defined the region-specific proteomic features of seven COVID-19 pulmonary structures by statistical analysis. Well-established cell type markers were identified in respective protein clusters, confirming the accuracy of our LMD experiments. Functional analysis demonstrated the enrichment of cell types and biological processes in each region with distinct features. Notably, we observed and validated the elevated expression of significant markers in several regions, such as macrophage marker MRC1 in ASI region, TGF-β pathway regulator TGFBI and EMT marker FN1 in PF region, as well as neutrophil marker MPO and NET marker Cit-H3 in BMP region. Interestingly, cathepsin CTSL and host cell receptor NPC1 were enriched in HAT2, highlighting the critical role of HAT2 in COVID-19 pathogenesis. Finally, we focused on the dysregulation of proteins related to viral entry, host restriction, and inflammatory response in multiple regions referring to a manually curated database. Dozens of key receptors and regulators were up-regulated compared to non-COVID-19 controls, including previously reported and predicted COVID-19 related regulators and potential therapeutic targets. We validated IFITM3, FURIN, HGF, and OSMR here, underlining their significance in further mechanistic research.

Collectively, we characterized the regional and spatially resolved proteome of COVID-19 pulmonary injuries in post-mortem tissue specimens with an unprecedented depth. A spectrum of protein and function dysregulations especially relevant to immune response were observed. This study offers unique insights for delineating the mechanisms of severe COVID-19 pulmonary injury and provides a valuable reference for identifying potential target for the therapeutic intervention.

### Limitations of the study

Compared to state-of-the-art single cell sequencing technologies, the current LMD-based workflow faces significant limitations in terms of spatial resolution and throughput, which ultimately restrict the data dimension of spatial proteomics. Further technical development is necessary to enhance the cutting precision and collecting scale of LMD platforms. On the other hand, the cohort in this study is relatively small in size, owing to the decrease in COVID-19 mortality rates at present. Future investigations should incorporate a larger number of specimens encompassing various subtypes of pneumonia.

## METHODS

### Ethics statement

COVID-19 lung samples of 11 individuals and non-COVID-19 lung samples of 3 individuals in this study were derived from autopsies from Wuhan Jinyintan Hospital and Tongji medical school, Huazhong University of Science and Technology, Wuhan, China. This study was approved by the ethics committees at Wuhan Jinyintan Hospital Ethics Committee (permission number: KY-2020-15.01) and Tongji medical school, Huazhong University of Science and Technology (permission number: [2021] IEC-A001). Written informed consent was obtained from the patients’ families with the permission for publication of all the research results. Laboratory confirmation of SARS-CoV-2 infection was performed at Wuhan Jinyintan Hospital and Wuhan Central Hospital, Wuhan, China. SARS-CoV-2 viral RNA was confirmed by real-time quantitative PCR or pathological staining.

### H&E staining, histology, immunohistochemistry, and immunofluorescence

The post-mortem tissue specimens were immediately fixed in 4% neutral formaldehyde for at least 72 h and embedded in paraffin as FFPE blocks stored at room temperature. For histological staining and spatial proteomic study, the tissues were cut into multiple sequential 5.0-µm sections with a microtome (RM2255, Leica). H&E staining were performed according to the standard procedures, and the results were reviewed by at least two histopathologists independently. For immunohistochemistry (IHC) and immunofluorescence (IF) staining, slides were deparaffinized and rehydrated, followed by 15 min of heat-induced antigen retrieval with EDTA (pH 9.0) in a microwave oven. The slides were washed with 0.02% Triton X-100 in phosphate-buffered saline (PBS) and then blocked with 5% bovine serum albumin (BSA) at room temperature for 1 h. The IHC staining was performed with a diaminobenzidine (DAB) detection kit (Dako). The multiplexed IF staining was performed following the instruction of Opal 7-color IHC kit (NEL811001KT, PerkinElmer). Other antibodies and staining reagents used in this study include rabbit anti-smooth muscle actin (SMA) antibody (1:200, 55135-1-AP, proteintech), mouse anti-TGFBI antibody (1:500, 60007-1-Ig, proteintech), mouse anti-fibronectin (FN1) antibody (1:400, GB12091, Servicebio), rabbit anti-TTF1 (NKX2-1) antibody (1:200, ab76013, Abcam), mouse anti-Cytokeratin 8 (KRT8) antibody (1:1000, GB12233, Servicebio), rabbit anti-human Myeloperoxidase (MPO) antibody (1:200, ab9535, Abcam), rabbit anti-H3-Cit antibody (1:500, ab5103, Abcam), mouse anti-human CD206 antibody (1:1000, ab300621, Abcam), rabbit anti-OSMR antibody (1:100, 10982-1-AP, proteintech), rabbit anti-IFITM3 antibody (1:800, 11714-1-AP, proteintech), rabbit anti-HGF antibody (1:200, A1193, ABclonal), rabbit anti-furin antibody (1:800, 18413-1-AP, proteintech) and DAPI (10236276001, Roche). The bright-field and immunofluorescent images were acquired using a Pannoramic MIDI system (3DHISTECH) and confocal microscopy (STELLARIS 8 WLL, Leica).

### Laser microdissection

The 5.0-µm FFPE sections for laser microdissection (LMD) were mounted onto PEN-membrane glass slides (Leica) following by deparaffinization, rehydration, H&E staining, and dehydration without covering slips. Stained tissue sections were scanned using a NanoZoomer S60 system (Hamamatsu) at 20× magnification prior to LMD. The obtained whole slide images were evaluated by pathological and cell type examination first and then subjected to cell type dissection and collection using a LMD7000 microscopy (Leica). The cutting parameters were optimized to minimized laser damage at first. Ten pathological regions with well-defined morphology characteristics were selected and dissected under the supervision of an experienced pathologist. For each sample, an accumulated area of 1 mm^2^ was collected into a 0.2 mL PCR tube for further processing.

### Proteomics sample preparation

The microdissected tissue samples were lysed and de-crosslinked in 50 μL lysis buffer containing 1% n-dodecyl β-D-maltoside (DDM), 600 mM guanidine HCl, 150 mM NaCl, and 10 mM HEPES (pH 7.4). After brief centrifugation, the samples were sonicated in a non-contacting BioRuptor Pico system (Diagenode) for 30 cycles at 4 °C (30 s-on and 30 s-off) and then heated at 95 °C for 90 min. The obtained tissue lysates were processed using a modified SISPROT protocol as we previously described.^19^ In brief, the tissue lysate was acidified to pH 3 by adding 1% formic acid (FA) before loaded onto an activated SISPROT tip packed with 0.6 mg mixed POROS beads of SAX and SCX (Applied Biosystems) and one plug of C18 disk (Empore). After cleaning by acetonitrile (ACN), the enriched proteins were reduced by 10 mM TCEP, alkylated by 40 mM chloroacetamide, and digested by 10 ng trypsin (Promega) at 37 °C for 60 min. The digested peptides were then transferred to the C18 layer by 500 mM NaCl, desalted by 1% FA, eluted to glass inserts by 80% ACN, 0.5% HOAc, and vacuum-dried in a SpeedVac machine (Thermo Fischer). To increase the depth of protein identification, part of the spectral library was built by in-solution digestion of protein lysate extracted from FFPE tissue sections and high-pH reversed-phase peptide fractionation as described previously.^19^ Briefly, FFPE sections from different blocks were dewaxed and collected into one centrifuge tube by scalpel. The tissue was then lysed and de-crosslinked in 1% SDS, 300 mM Tris-HCl (pH 8.0) by sonification and then heating at 95 °C for 60 min. One milligram of proteins was extracted, reduced, alkylated, and digested using a modified in-solution sample preparation protocol.^94^ One hundred micrograms of desalted peptides were fractioned on an microflow HPLC (1260, Agilent) using a XBridge peptide BEH C18 column (130 Å, 5 μm, 2.1 mm × 150 mm) within 60 min linear gradient from 2% to 30% ACN in 10 mM ammonium bicarbonate (pH 8.0) at a flow rate of 0.2 mL/min and concatenated into 24 fractions. The fractions were then desalted using homemade C18 tips and vacuum-dried. All peptide samples were re-dissolved in 0.1% FA with spiked iRT peptides (Biognosys). For LMD samples, each sample was injected twice for ddaPASEF and diaPASEF analysis, respectively.

### LC-MS analysis

All LC-MS/MS analysis were performed on a timsTOF Pro (Bruker Daltonik) mass spectrometer coupled to a nanoElute UPLC system (Bruker Daltonik) *via* a captive spray ion source. Peptides were loaded and separated on a 50 μm i.d. × 20 cm capillary column in-house packed with C18 beads (1.9 μm, 120 Å, Dr. Maisch GmbH) at a flow rate of 100 nL/min and the column temperature was kept at 50 °C in a column oven. For all samples, the peptides were separated using a linear gradient from 2% to 22% phase B in 50 min and stepped up to 35% in 10 min followed by a 20 min wash at 80% phase B where phase A was 0.1% FA in ddH_2_O and phase B was 0.1% FA in ACN. For ddaPASEF analysis, parameters were set as we previously described.^20^ Precursor ions in m/z range of 300 to 1500 were scanned in positive electrospray mode. The ion mobility range was scanned from 1/K_0_ = 0.75 to 1.30 Vs/cm^2^. The ramp time was 200 ms and the total cycle time was 1.03 s which comprised 1 MS1 survey tims-MS and 4 PASEF MS/MS scans per acquisition cycle. Only doubly and triply charged features were selected to trigger MS/MS scans. For diaPASEF analysis, 32 isolation windows of 20 m/z width were defined with m/z range from 400 to 1040 and mobility range from 1/K0 = 0.75 to 1.30 Vs/cm^2^. The collision energy was set to rise linearly over the covered mobility range from 59 eV at 1/K0 = 1.6 Vs/cm^2^ to 20 eV at 1/K0 = 0.6 Vs/cm^2^. Total resulting DIA cycle time was estimated to be 1.85 s.

### MS data analysis

All MS raw files were searched by the Spectronaut software (version 16.2, Biognosys). For ddaPASEF-based spectral library generation, we utilized MS files from 24 peptide fractionation samples and 71 second injection of LMD samples. The combined raw files were searched against a combined Homo sapiens (UP000005640, 20,593 entries, downloaded on 1st Oct, 2022) and SARS-CoV-2 (UP000464024, 17 entries, downloaded on 1st Oct, 2022) UniProt fasta database using the Pulsar engine embedded in Spectronaut with all parameters set as default. The resulting library comprised 210,597 precursors accounting for 168,887 peptides and 11,438 protein groups. Next, a default library-based analysis pipeline was applied for protein identification and quantification of all DIA raw files.

### Statistical analysis

Statistical analysis and visualization of proteome were performed using Perseus software (2.0.7.0) and R studio (4.1.0). After excluding potential contaminants manually, we filtered proteins identified in this study for at least two valid values in at least one region. Missing values were imputed by the strategy of one sample’s normal distribution of proteome abundance using Perseus (down-shifted mean by 1.8 standard deviation and scaled by 0.3). One-way ANOVA and student’s t test were performed for regional enriched analysis and paired group comparison, respectively. In regional enriched protein cluster analysis, log_2_ fold change (FC) was calculated by the median of protein expression ratio between one region and the rest, adjusted p values were calculated using Benjamini–Hochberg (B-H) method, proteins with FC larger than 1.2 and adjusted p value smaller than 0.05 were kept for following analysis. The clusterProfiler package (4.0.5) in R and an online platform Metascape^51^ were used for GOBP and KEGG enrichment analysis and visualization. To construct a COVID-19 related functional protein database, we manually integrated databases of COVID-19 pathogenesis related proteins, including virus receptor, viral entry-cofactor, viral-entry related protease, and inflammatory related proteins, based on reported and predicted datasets, which contained 273 proteins. ^13, 79–83^ The significant up-regulated proteins were identified using LIMMA with p-value < 0.05 and Fold Change > 2.

## DATA AVAILABILITY

The MS raw files in this study have been deposited in the ProteomeXchange Consortium *via* the PRIDE partner repository. All other data that support the findings of this study are provided in the Supplementary Information and Source Data files. Source data are provided with this paper.

## ACKNOWLEDGEMENTS

This work was supported by the China State Key Basic Research Program Grants (2020YFE0202200, 2021YFA1301601, 2021YFA1301602, 2021YFA1302603, and 2022YFC3401104), the National Natural Science Foundation of China (22074060, 22125403, 92253304, and 22150610470), the Shenzhen Innovation of Science and Technology Commission (JCYJ20200109140814408, JCYJ20200109141212325, and JCYJ20210324120210029), the National Key R&D Program of China (2021YFC2300901), the Coronavirus Pneumonia Project from the Ministry of Science and Technology of China (2020YFC0844700), and the Natural Science Foundation of Hubei Province (2021CFA053). We thank Juan Min and Ding Gao from Center for Instrumental Analysis and Metrology of the Wuhan Institute of Virology for their technical support.

## AUTHOR CONTRIBUTIONS

Z.L.S., R.T., and Y.Z. conceived and supervised this project. Y.Z., L.M., L.L., Q.L., and L.R. participated in autopsy sample collection and histopathological examination. Y.C., Y.M., and X.W. performed the laser microdissection experiments. Y.M., X.W., and Y.X. performed the proteomic experiments. L.Y., A.H., W.G., Y.M. and Y.C. participated in data analysis and visualization. C.Y. performed IHC and IF experiments. Y.M., Y.C. Y.L., Y.Z., R.T., and Z.L.S. wrote and edited the manuscript with inputs from co-authors.

## CONFLIT OF INTERESTS

The authors declare no conflict of interest.

