## Supplementary material for "Deep spatial proteomic exploration of severe COVID-19-related pulmonary injury in post-mortem specimens": Figure S

A

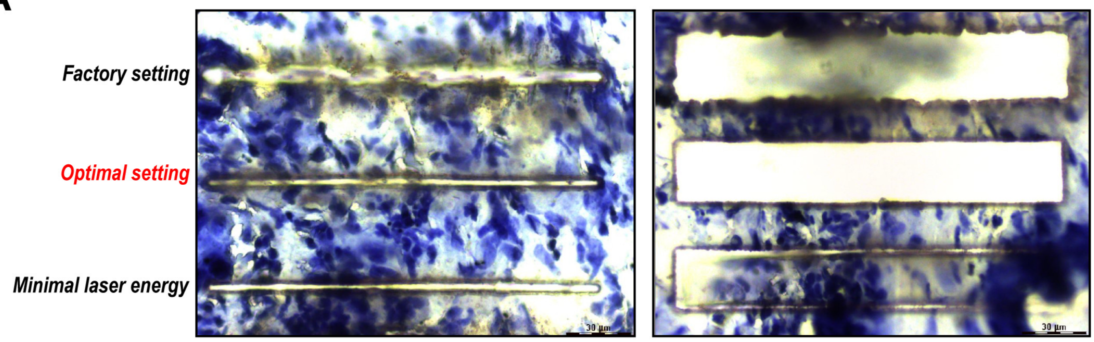


B

| | Power | Aperture | Speed | Head current | Pulse frequency |
|---|---|---|---|---|---|
| *Factory setting* | 35 | 15 | 10 | 80% | 120 |
| *Optimal setting* | 5 | 1 | 20 | 100% | 240 |
| *Minimal laser energy* | 1 | 1 | 20 | 80% | 120 |

**Figure S1. Laser parameter optimization.**

(A) Representative images of tissue section after laser microdissection using parameters of factory setting, minimal laser energy, and optimal setting, respectively.

(B) Key parameters of three different laser settings.

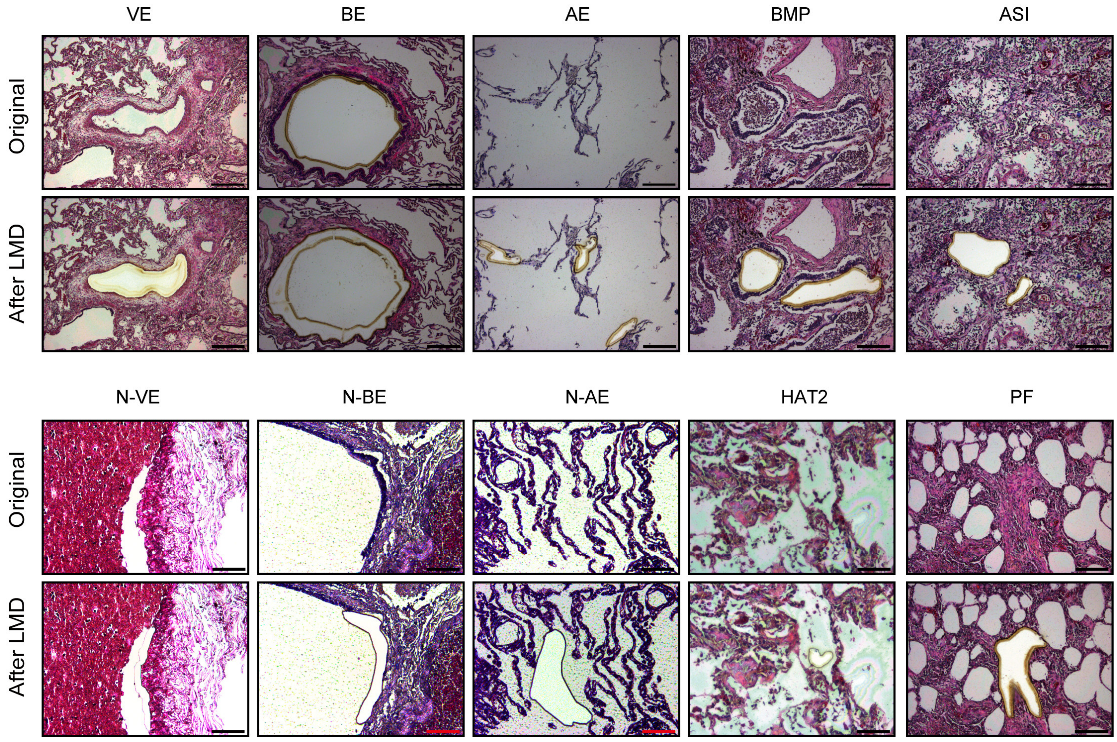


**Figure S2. Representative H&E staining images of ten pulmonary regions before and after laser microdissection, respectively.**

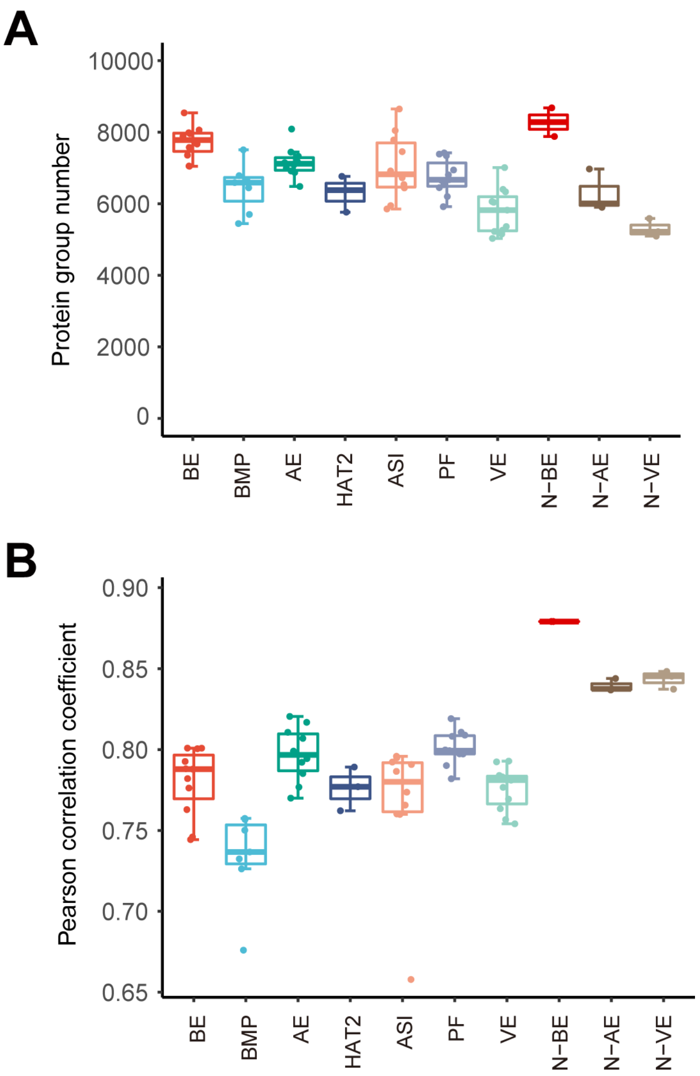


**Figure S3. Proteome depth and intra group reproducibility.**

(A) Box-whisker plot of identified protein number in ten dissected regions, respectively.

(B) Box-whisker plot of Pearson correlation coefficient in ten dissected regions, respectively.

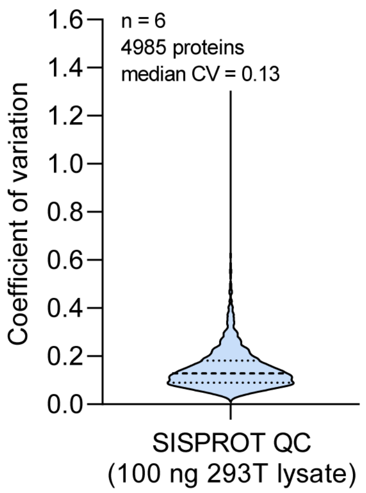


**Figure S4. Violin plot showing the distribution of coefficient of variation (CV) values in SISPROT quality control samples (100 ng 293T cell lysate per sample with six technical replicates).**

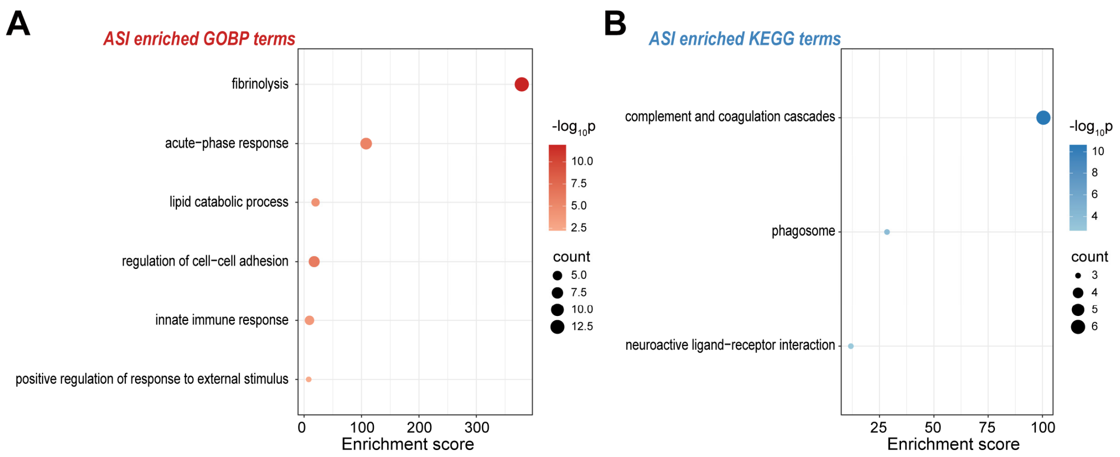


**Figure S5. Proteomics-based functional analysis of airspace inflammation in COVID-19 lungs.**

(A) Dot plot showing all significant GOBP terms for ASI-enriched protein cluster. The size of circle represents the number of proteins and the color indicates the $-\log_{10}$ p-value.

(B) Dot plot showing all significant KEGG terms for ASI-enriched protein cluster. The size of circle represents the number of proteins and the color indicates the $-\log_{10}$ p-value.

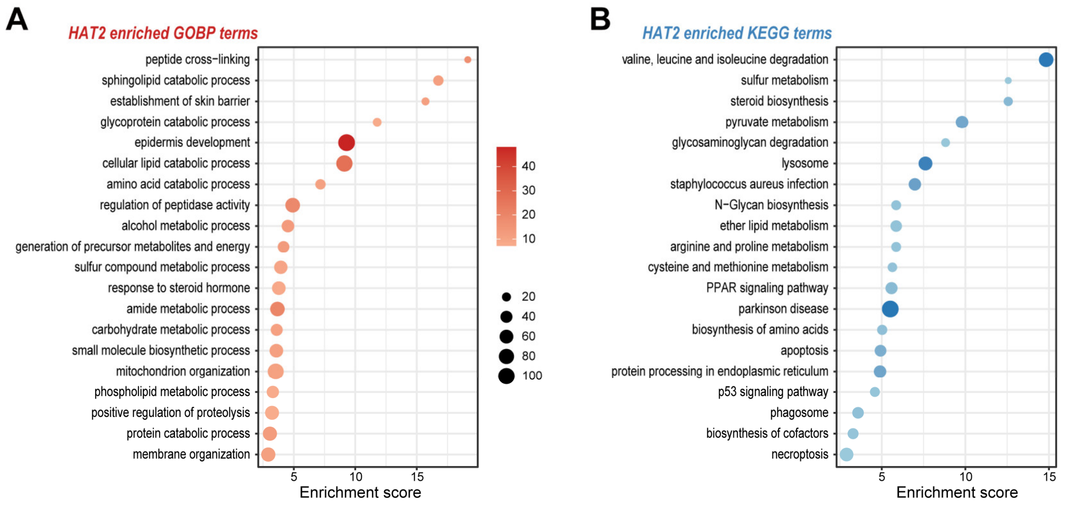


**Figure S6. Proteomics-based functional analysis of hyperplastic alveolar type 2 in COVID-19 lungs.**

(A) Dot plot showing top-20 significant GOBP terms for HAT2-enriched protein cluster. The size of circle represents the number of proteins and the color indicates the -log10 p-value.

(B) Dot plot showing top-20 significant KEGG terms for HAT2-enriched protein cluster. The size of circle represents the number of proteins and the color indicates the -log10 p-value.
